# Selection Of Models For The Regression Problems In Biology Using Chi-Square Test

**DOI:** 10.1101/2022.09.08.507150

**Authors:** Aaditya Prasad Gupta

## Abstract

Biological systems, at all scales of organization from nucleic acids to ecosystems, are inherently complex and variable. Therefore mathematical models have become an essential tool in systems biology, linking the behavior of a system to the interaction between its components. Parameters in empirical mathematical models for biology must be determined using experimental data, a process called regression because the experimental data are noisy and incomplete. The term “regression” dates back to Galton’s studies in the 1890s. Considering all this, biologists, therefore, use statistical analysis to detect signals from the system noise. Statistical analysis is at the core of most modern biology and many biological hypotheses, even deceptively. Regression analysis is used to demonstrate association among the variables believed to be biologically related and fit the model to give the best model. There are two types of regression, linear and nonlinear regression to determine the best fit of the model. In this manuscript, we perform a least squares error fit to different models and select the best fit model using the *χ*^2^-test, and determine the p-value of the selected model to data that was collected when various doses of a drug were injected into three animals, and the change in blood pressure for each animal was recorded.

## 1 Introduction

A model is a mathematical description of a physical, chemical, or biological state or process whose part are mathematical concepts such as constant, variables, function, equation, inequalities, etc. Using a model help us to think about such processes and their mechanisms, so we can design better experiments and comprehend the results. Models are tools that assist in theory development^1^. Criteria for selecting a model include the goodness of fit, freedom from systematic error and simplicity^2^. To fit the model we need regression. The regression model is an equation that defines the outcome, or dependent variable Y, as a function of an independent variable X, and one or more model parameters. The regression model determines a relationship between an independent variable and a dependent variable by providing a function. Regression analysis is often used to demonstrate association among variables believed to be biologically related. Linear and non-linear regression fit a mathematical model to the data to determine the best values of the parameters of the model.

For this, we need to select the best model to obtain our maximum output result^3^. Model selection is the task of selecting a statistical mode from a set of candidate models, given data. There are different criteria for the selection of data such as Final Prediction Error (FPE), Alkaike’s Information Criterion (AIC), F-test, Shortest Data Descriptor (SDD), Residuals, Parameter Variance Statistics, Determinant Comparison, Zero Pole-Zero Cancellation, etc. Applying these criteria we have a different test to select the best model. In this manuscript, we perform the chi-square test for the goodness of fit. However, there are other tests such as the Exact Test of goodness-of-fit, Power analysis, G-test of goodness-of-fit, Chi-square test for independence, G-test of independence, Fisher’s exact test, Small numbers in chi-square and G-tests, Repeated G-tests of goodness-of-fit and Cochran-Mantel-Haenszel test, etc but we use Chi-square test of goodness-of-fit for our manuscript. The Chi-square goodness of fit test is a test that determines whether the variable is likely to come from a specified distribution or not. It was proposed by Pearson (1900)^4^. It is also known as statistical tests^5^. Modeling has been widely used in biological research. This manuscript is arranged to incorporate the information about model fitting in biology. In section 2 we discuss a short description of regression in biology. Similarly, in section 3 we discuss the data used in this manuscript whereas in section 4 we discuss a different way of obtaining parameters and uncertainties for fit in python. While section 5 discusses the result. Finally, this manuscript ends with a discussion and a brief conclusion in section 6.

## 2 Regression Problem in Biology

Regression involves the determination of the degree of relationship in the patterns of variation of two or more variables through the calculation of the coefficient of correlation. The regression problem is how to model one or several dependent variables/responses, y, using a set of predictor models. It typically corresponds to very different biological questions. Evolution regression across species is one of the major statistical procedures used to study the evolutionary relationship between biological variables and test hypotheses^6^. In biology, compartmental analysis models are used in radio-active tracers to study metabolic disorders^7^. Over the last decades, sophisticated statistical models have been developed. Most of these developments have been focused on solving statistical problems and biological interpretations. assumptions made to solve statistical problems are often incompatible with biological processes. So both biological and statistical considerations are necessary to obtain meaningful evolutionary and allometric regression model observation or measurement error is a serious concern for most comparative studies^8^..

## 3 Data

The dataset for this model is retrieved from a book named “Fitting Models to Biological Data using Linear and Nonlinear Regression”^9^. In this model, various doses of a drug were injected into three animals, and the change in blood pressure for each animal was recorded. The data for this model is fitted with nonlinear regression. The dataset includes 2 variables along with 4 parameters. The variables used in this dataset are shown below in table 1.

**Table 1.** Different doses in three different animals

| log(dose) | $Y_1$ | $Y_2$ | $Y_3$ |
| --- | --- | --- | --- |
| -7.0 | 1 | 4 | 5 |
| -6.5 | 5 | 4 | 8 |
| -6.0 | 14 | 14 | 12 |
| -5.5 | 19 | 14 | 22 |
| -5.0 | 23 | 24 | 21 |
| -4.5 | 26 | 24 | 24 |
| -4.0 | 26 | 25 | 24 |
| -3.5 | 27 | 31 | 26 |
| -3.0 | 27 | 29 | 25 |

In the above table 1, log (doses) denotes the logarithm of doses of drug given to the animals whereas *Y*_1_, *Y*_2_, and *Y*_3_ are the change in blood pressure to whom various doses were injected. *Y*_1_ is the value for the minimal curve asymptote (theoretically, the level of response, if any, in the absence of a drug), *Y*_3_ is the value for the maximal curve asymptote (theoretically, the level of response produced by an infinitely high concentration of drug), and *Y*_2_ is the value produces the response halfway between the Top and Bottom response levels (commonly used as a measure of drugs’ potency)

## 4 Fit in Python

Python has methods for finding a relationship between data points and drawing a regression function that can be both linear and non-linear. In statistics, we say that regression is linear when it’s linear in the parameters. Fitting linear models is an easy task, we can use the least squares method and obtain the optimal parameters for our model. However, not all problems can be solved with pure linear models. So nonlinear model has required that guarantee useful mathematical properties and are also highly interpretable. A nonlinear model is a mathematical model that fits an equation to certain data using a generated line such as a curve that is not a straight line. The regression problems are generally solved using curve fit and machine learning methods. Curve fitting examines the relationship between one or more predictors (independent variables) and a response variable (dependent variable), intending to define a “best fit” model of the relationship. Using the curve fitting makes our outcomes much more interpretable and even guarantees some mathematical property on our model output. Here, to perform the regression, we use the curve fit methods. Besides these, we use Matplotlib and Numpy. These are the main libraries we’ll use for fitting purposes.

In python, the curve fit can be formed using several modules such as Scipy and lmfit based on least-squares minimization. A common use of least-squares minimization is curve fitting, where one has a parametrized model function meant to explain some phenomena and wants to adjust the numerical values for the model so that it most closely matches some data. Here, we make rigorous use of Scipy module from the Python. Scipy stands for Scientific Python. It is the scientific computing module of Python providing in-built functions on a lot of well-known Mathematical functions. With scipy, curve fitting problems are typically solved with scipy.optimize.curve_fit, which is a wrapper around scipy.optimize.leastsq. Like scipy.optimize.curve_fit, lmft uses a function that is meant to calculate a model for some phenomenon, and then uses that to best match an array of supplied data. In curve fitting, the uncertainties of the fitted parameters are a measure of how strongly the parameter depends on the data.

### 4.1 Least Squares Fitting

The method of least squares is a standard approach in regression analysis to approximate the solution of over-determined systems (sets of equations in which there are more equations than unknowns) by minimizing the sum of the squares of the residuals (a residual being the difference between an observed value and the fitted value provided by a model) made in the results of each equation.

Let *y*_*i*_ be the observed set of n data points, and *f* (*x*_*i*_, *a*_1_, *a*_2_, …, *a*_*n*_) be the function that is fitted to the data set. The square of the reside (*R*^2^) is calculated as

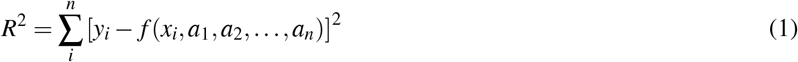

The *R*^2^ is minimized and the condition for *R*^2^ to be a minimum is that

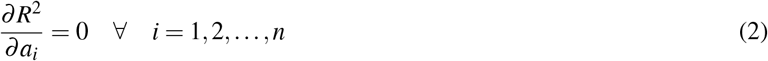

For a linear fit,

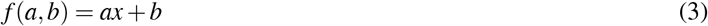

Then,

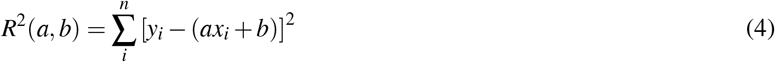

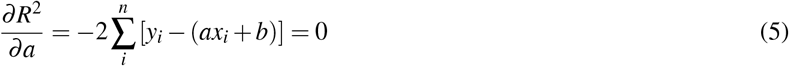

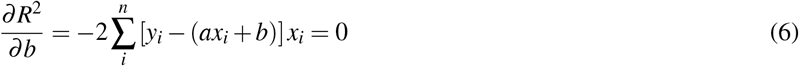

These lead to the equations

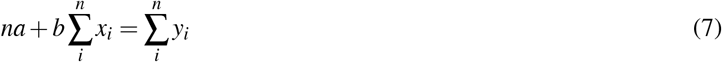

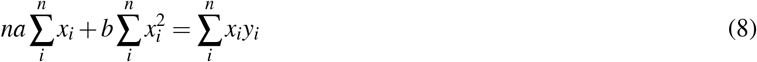

In the matrix form:

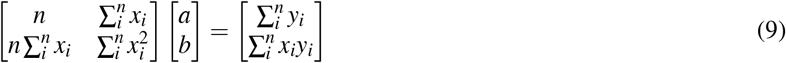

Thus

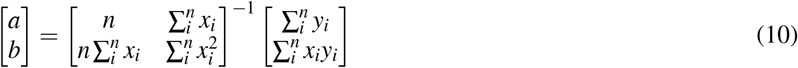

Similarly

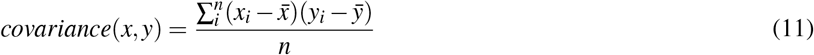

## 5 Results

In regression analysis, curve fitting is the process of specifying the model that provides the best fit to the specific curve in our dataset. Here, we tried the following models represented by equation 12, 13, 14 and 15 in our fit. Equation 12 is a linear equation in independent variable x as function of dose and dependent variable y as response. In this model, we have a and b as the parameters. Similarly, equation 13 is a quadratic equation that defines the response (dependent variable, y) and function of dose (independent variable, x), with the model parameters a, b and c. Equation 14 is a cubic equation in x and y defined as in equation 12 and equation 13, with the model parameters a, b, c, and d.

The 15 shows a standard model named as Hill function. It is used to model density-dependent growth or decline. Here it is used to model a process when various doses of a drug are applied to animals and the corresponding response was measured. The response (also called the independent variable, y) is measured as a function of dose (also called the independent variable, x). The model parameters are Bottom, which denotes the value of y for the minimal curve asymptote (theoretically, the level of response, if any, in the absence of drug, and Top which denotes the value of y for the maximal curve asymptote (theoretically, the level of response produced by an infinitely high concentration of drug), LogEC50 which denotes the logarithm of drug dose (or concentration) that produces the response halfway between the Top and Bottom response levels (commonly used as a measure of drugs’ potency), and the Hillslope which denotes the steepness of the drug-response curve (often used as a measure of the sensitivity of the system to increments in drug concentration or doses.

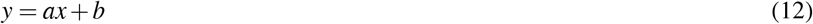

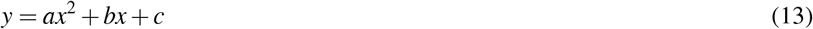

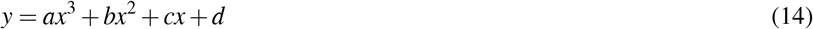

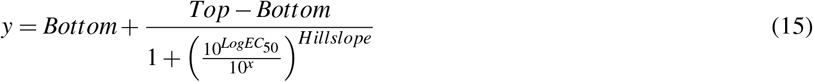

Curved relationships between variables are not as straightforward to fit and interpret. In order to fit in the model represented by equations 12, 13, and 14, we assume bottom as lower error (Δ*L*), and Top as upper error (Δ*U*). These two error are asymmetric, and they are made symmetric using error 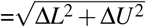. Then the data that is fitted to the model using Scipy and the fitted parameters are determined.

Figure 1 and figure 2 show the data with asymmetric error and symmetric error respectively. Along the x-axis we have a log of the Dose that is applied to the animals and along the y-axis, we have a change in the blood pressure of animals along the y-axis while doses are given to them. Uncertainty has of great importance to avoid biased results. The asymmetric errors are converted to symmetric ones because the least squares algorithm like curvefit will not be able to calculate the loss function negative error.

**Figure 1.**
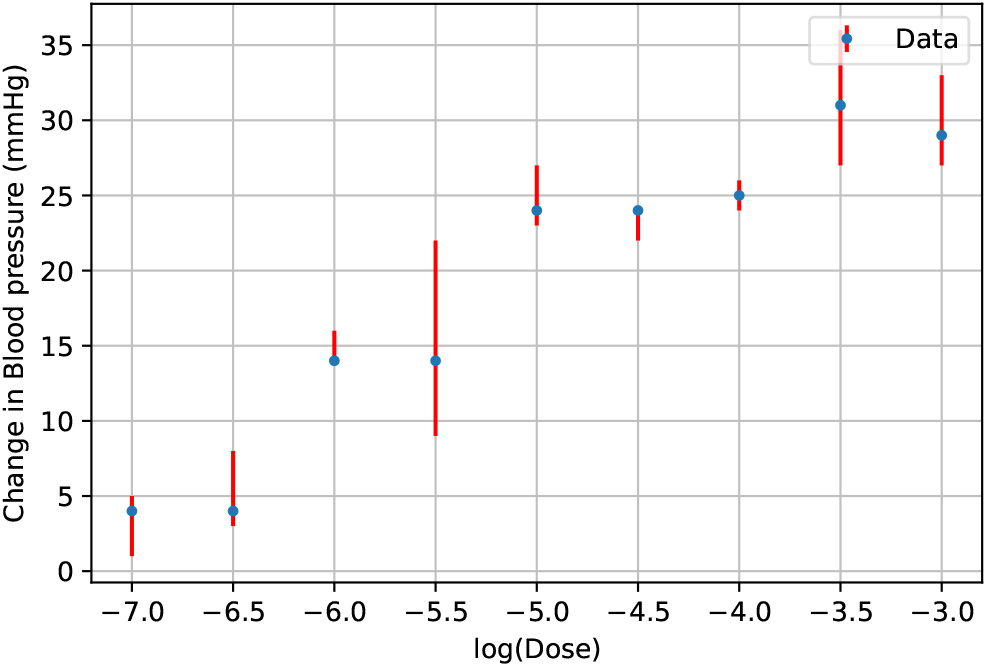
Data plot with asymmetric error

**Figure 2.**
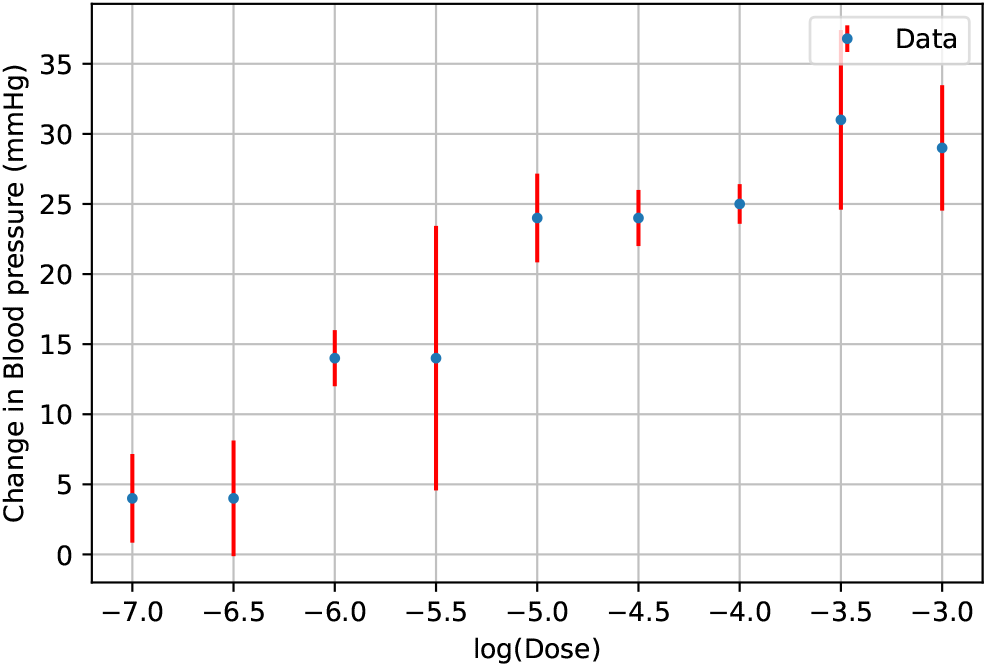
Data plot with symmetric error

The figures 3, 4, 5 and 6 show the plot for both data along with uncertainties and the corresponding model that is fitted to it. Along the x-axis we have a log of the Dose that is applied to the animals and along the y-axis, we have a change in the blood pressure of animals along the y-axis while doses are given to them. Figure 3 shows the straight line fit to the data, while figure 4 shows the quadratic fit. Similarly, figure 5 and figure 6 show the cubic equation fit and Hill function fit respectively. The fitted data formed a curve that is fitting the maximum data and providing us with the best fit for our data. Figure 6 does not have uncertainties in the response value as the Hill function have the parameters to accommodate the lower and the upper values of the change in blood pressures.

**Figure 3.**
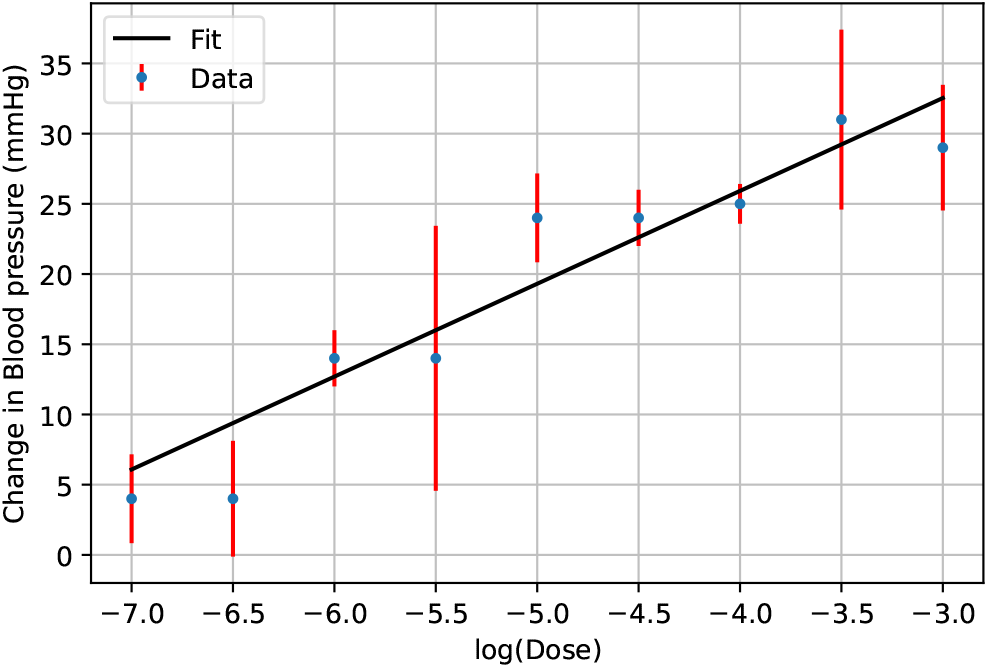
Fit plot with data for linear model

**Figure 4.**
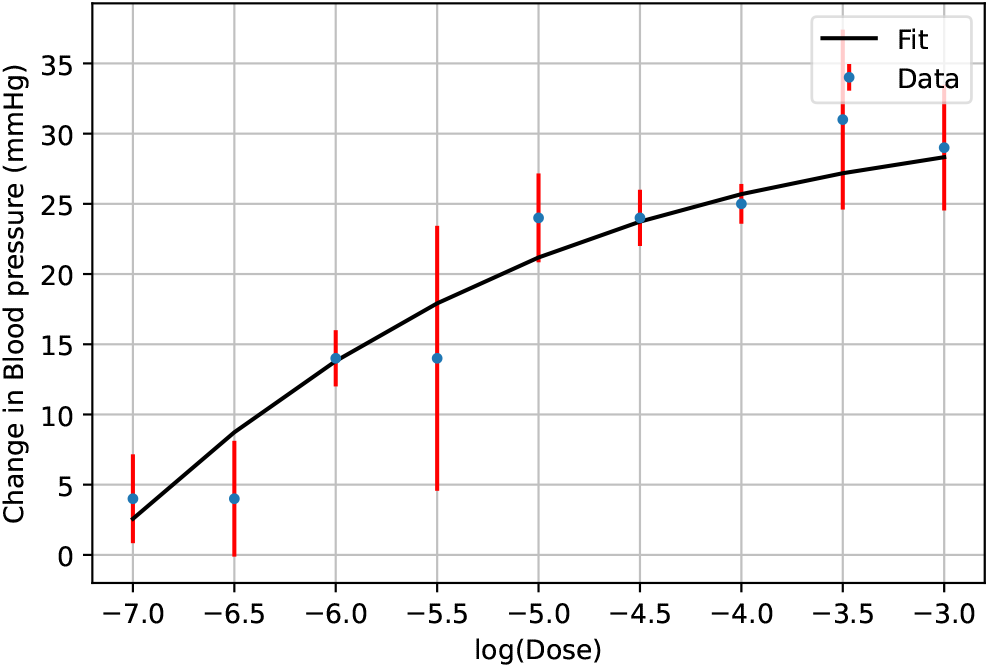
Fit plot with data for quadratic model

**Figure 5.**
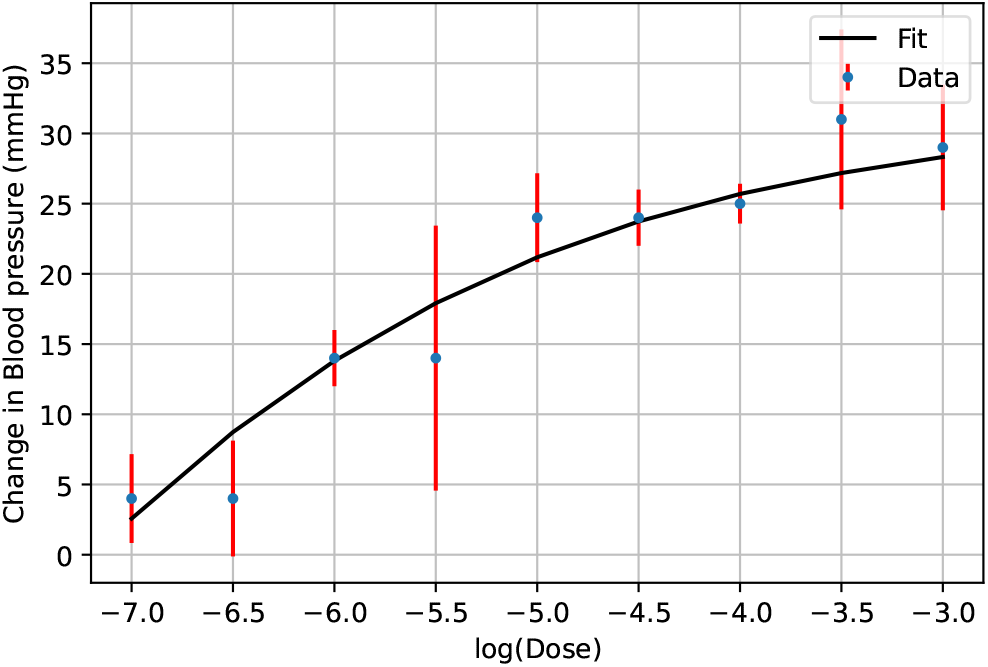
Fit plot with data for cubic model

**Figure 6.**
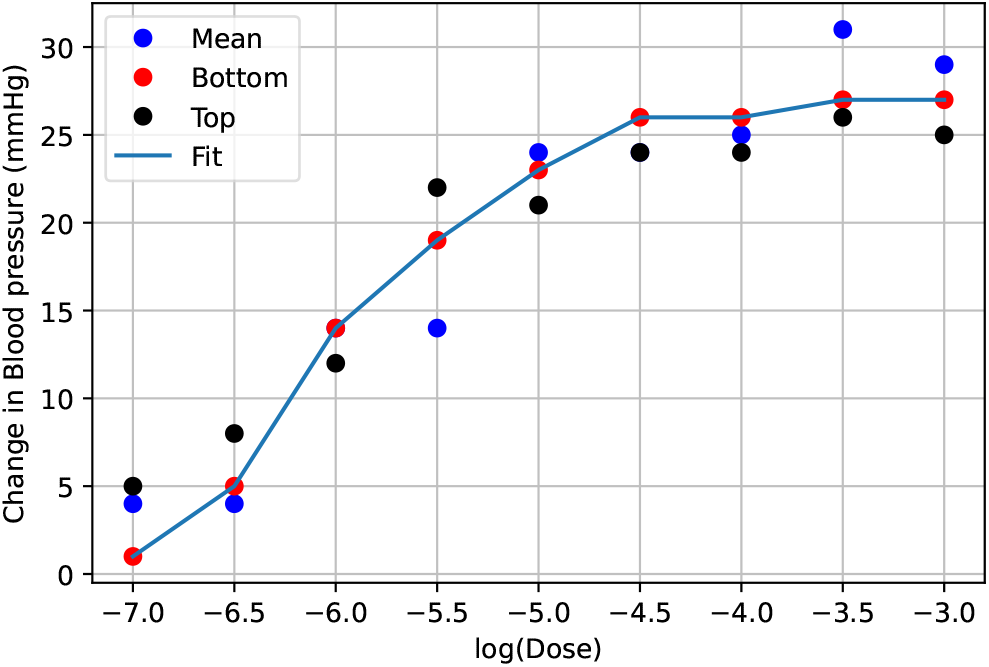
Fit plot with data for Hill function model

Lastly, the table 2 shows the chi-square value, degree of freedom (dof), and reduced chi-square value of the different models which is essential for this manuscript. The Chi-square test is a statistical procedure for determining the difference between the observed and expected test. It is very important to study the chi-square test of the models because it allows us to assess the trained regression model’s goodness of fit on the training, validation, and test datasets. Reduced chi-square is a chi-square per degree of freedom used extensively in the goodness of fit testing. The expected value for the best model is 1 or more near 1. The model has 1 value indicates the difference between observed data and fitted data has a similar magnitude of weight. The degree of freedom (dof) is the maximum number of logically independent values which are values that have the freedom to vary in the data sample. Table 2 shows that for Hill function, it has a chi-square value of 11.4923, dof value of 7 and the reduced chi-square value of 1.6417. Similarly, for the linear model, the chi-square value is 6.4188, the dof is 7 and the reduced chi-square is 0.9169 which indicates that the model fits the best compared to the Hill function as it is near 1. While the reduced chi-square value for the quadratic model and cubic model are 0.1666 and 0.2000 which are quite far from 1. The Hill function has a p-value of 0.1185, while the p-value for the linear model is 0.4918.

**Table 2.**
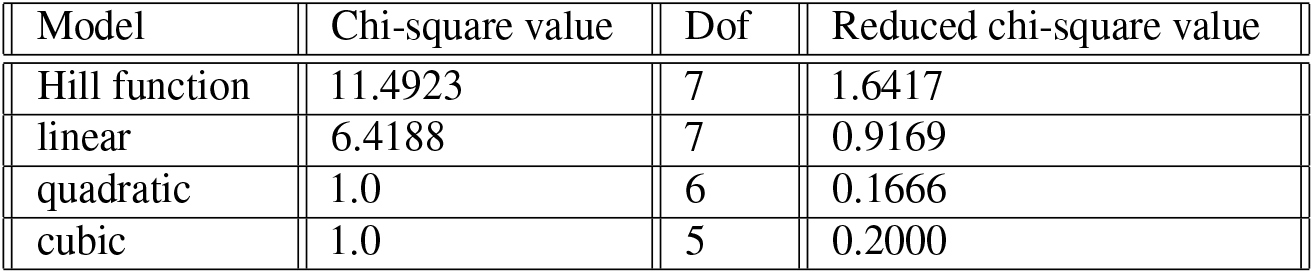
Different models with their chi-square value

## 6 Discussion and Conclusions

In this manuscript, different models were fitted to the data, and the chi-square test of the respective model was performed. Python modules were used which have many excellent and wee-maintained libraries that facilitate high-level scientific computing analyses. The module such as numpy^10^ offer a numerical processing library that supports multi-dimensional arrays, matplotlib^11^ which offers a plotting library with tools to display data in a variety of ways and SciPy^12^ allows for minimizing or maximizing objective function, possibly subject to constraints and find the optimized parameter set for a stated function that perfectly fits the provided data set were used in Anaconda^13^.

We found that regression analysis, especially nonlinear regression, is an essential tool to analyze biological (and other) data. The regression in the biological system sometimes becomes a failure to demonstrate a statistically significant unbiased relationship due to many factors. In this manuscript, we had data on the response of doses of drugs in different animals. In the Hill function, the data were fitted to the standard model considering the different parameters to accommodate the lower and the upper values of the change of blood pressures without error. The data were fitted in different models such as linear, quadratic, and cubic with the parameter and having uncertainty. The uncertainty in the data has great importance to avoid biased. When the uncertainties are not small, asymmetric, and abnormally distributed, one needs more sophisticated ways, which we have done in our manuscript. The Hill function and linear model have reduced chi-square values near the ideal value, while those for quadratic and cubic are too small and considered for further study. The Hill function has a p-value of 0.1185, while the p-value for the linear model is 0.4918. Thus, they cannot be rejected at the 90% confidence interval.

